# Genomic insights into antiviral defense systems in haloarchaea and their impact on virus susceptibility

**DOI:** 10.1101/2025.11.08.687337

**Authors:** Zaloa Aguirre-Sourrouille, Thomas Hackl, Hanna M. Oksanen, Tessa E.F. Quax

**Author notes:** Address correspondence to Tessa E.F. Quax.

## Abstract

The ongoing evolutionary arm race between archaea and their viruses has led to the development of diverse defense systems against viruses. While recent computational approaches have uncovered many bacterial antiviral defense systems, the viral infection strategies and antiviral responses of archaea remain poorly explored. In this study, we identified antiviral defense systems encoded in the genomes of 20 recently sequenced haloarchaeal strains. These systems were found to be representative of the broader repertoire of defense systems present across all 253 complete sequenced Halobacteria (class) genomes in the RefSeq database. Detailed analysis showed that these haloarchaea usually harbor multiple different defense systems, with a particularly high abundance of uncharacterized predicted defense systems against viruses (Phage Defense Candidates (PDCs)). To further explore the impact of anti-viral defense mechanisms on host range, an extensive virus-host pair screening was performed using a panel of known virulent viruses. By correlating the genomic defense profiles with observed viral infectivity and adsorption profiles, a weak correlation between the number of encoded defense systems and viral susceptibility was detected. Specifically, hosts infected by fewer viruses tended to encode a broader repertoire of antiviral defense systems, whereas those with fewer defense systems were frequently infected. It was found that the host range of haloarchaeal viruses is majorly determined by the availability of viral receptors, whereas the presence of anti-viral defense systems plays a smaller but significant role. These findings offer valuable insights into the evolutionary pressure shaping archaeal antiviral strategies and lay the groundwork for future functional studies of archaeal defense systems.

**Importance:** Archaeal viruses are understudied, compared to viruses infecting bacteria and eukaryotes. Haloarchaea have evolved as model organisms for the study of virus-host interaction in archaea, as they harbor the highest number of isolated archaeal viruses. Salt-loving haloarchaea dominate microbial communities in hypersaline environments, where they face constant viral threats. To survive, they have evolved a diverse array of antiviral defense systems. Until now, the relevance of these defense systems for susceptibility to viruses had not been tested in archaea. The findings of this study show that strains with a broader repertoire of defense systems tend to be less susceptible to viral infection, while those with fewer systems are more frequently targeted, suggesting an advantage for the accumulation of different defense systems. In addition, it was found that the major determinant of host range is the availability of viral receptors on the host cell. These insights provide a foundation for future research into the molecular mechanisms and ecological roles of archaeal defense systems.

## Introduction

Viruses are the most abundant biological entities in the biosphere [1] and infect members of the three domains of life. They are present in a great variety of environments and play important roles in the global biogeochemical cycles and dynamics of microbial populations [2]. Viruses infecting archaea show a remarkable morphological diversity and an impressive genetic repertoire, as 75% of their genes lack significant sequence homology to other known genes [3]. Despite their significant role in ecosystems, little is known about virus-host interactions in archaea.

In their natural environments, microorganisms face the constant threat of virus predation and therefore have evolved a diverse array of defense systems, each targeting different stages of the viral life cycle [4–6]. Consequently, viruses have developed complex counter-defense strategies to overcome these defenses [7, 8], emphasizing the selective pressure that drives the arms race between microorganisms and their viruses. This arms race is marked by high mutation rates and horizontal gene transfer, resulting in the rapid evolution of genetic traits and increased genetic diversity of hosts as well as of viruses [7, 9]. Moreover, this arms race is thought to have an impact on global nutrient cycling and climate [10, 11], on the dynamics of the biosphere [12] and on the evolution of virulence in human pathogens [13].

For a successful infection, viruses must first adsorb to the cell surface through recognition of specific host receptors. In bacteria, these receptors are often surface components [14]. To counteract viral infection, bacteria can alter their cell surface by random mutations or phenotypic variations, thus masking the receptors and leading to a decreased phage adsorption [15–19]. In archaea, however, only a limited number of receptors have been identified. Previous research has shown that archaeal viruses bind to the S-layer glycoproteins or to filamentous structures at the host surface [20–29]. Like bacteria, archaea can mutate their surface components in order to escape viral infection [24, 27, 30–33].

Once the viruses have entered the cell, their genomes are replicated before new viral particles are formed and released from the host cells. However, for viruses to be able to produce progeny virions, they must overcome several defense systems that bacteria and archaea express. Adsorption, viral entry, and intracellular defense mechanisms, influence the viral host range, which is determined as the spectrum of cell types and host species that a virus can infect and successfully produce progeny from [34]. The host range varies between different viruses. Recent studies in bacteria have demonstrated that the presence of antiviral defense systems correlates with host susceptibility to viral infection, whereby an increased number of such systems is associated with enhanced resistance to viruses [35]. Defense systems of bacteria and archaea can roughly be divided into (i) systems targeting viral nucleic acids (e.g., restriction-modification (RM) systems and clustered regularly interspaced short palindromic repeats and their associated Cas proteins (CRISPR-Cas)), (ii) abortive infection (Abi) systems that lead to the death of the infected host and (iii) other types of systems [4].

RM systems have been found in over 90% of all sequenced bacterial and archaeal genomes [36] and confer resistance to a wide variety of extrachromosomal elements such as viruses and plasmids [37]. In archaea, RM systems have been identified and described in thermophilic archaea [38–43], methanogenic archaea [44–46] and halophilic archaea [47,48]. CRISPR-Cas systems are found in around 90% of archaeal genomes and, frequently, multiple different CRISPR-Cas systems are present in a single genome [49, 50]. The first archaeal system in which CRISPR-Cas mediated defense was demonstrated *in vivo* was for members of *Thermoproteota*, in particular the strains *Saccharolobus solfataricus* P2 and *Saccharolobus islandicus* REY15A [51, 52]. Further research into the distribution of CRISPR spacers of acidothermophiles revealed matches to fuselloviruses, rudiviruses, and β-lipothrixviruses [53]. CRISPR-Cas systems have also been found in haloarchaea [54].

Abortive infection systems reported for archaea encompass toxin-antitoxin (TA) systems [59, 70, 71] and cyclic oligonucleotide-based antiphage signaling system (CBASS) utilizing abortive infection [72].

Other known defense systems found in archaea include superinfection exclusion [25, 55–59], dormancy [60], DNA phosphorothioation [61] and argonaute proteins [62–69]. Many bacterial defense systems have been found in archaea, but recent studies have also shown defense systems unique to archaea [73, 74]. Moreover, archaeal genomes contain many uncharacterized genes that might encode yet-to-be-discovered antiviral defense proteins and pathways. Although defense systems have been identified in archaea, it remains unclear how these systems influence viral host range or whether the accumulation of such defense systems makes the host more resistant to viral infections.

In this study, the first systematic analysis of the relation between the viral host range and antiviral defense systems encoded in the genomes of haloarchaeal hosts is presented. A panel of haloarchaea and their virulent viruses was used, taking advantage of the viruses’ ability to form plaques on the archaeal culture lawns. The defense systems present in a custom panel of 20 recently sequenced haloarchaeal strains were identified using bioinformatic tools and correlated to their susceptibility to several archaeal tailed viruses (arTVs) [75]. This analysis revealed a weak correlation between the number of antiviral defense systems and the viral susceptibility of the host strains. It also indicated that the availability of viral receptors on the host cell is a major determinant of the viral host range.

## Results

### Haloarchaea encode multiple defense systems

To date, most archaeal viruses isolated from hypersaline environments infect extremely halophilic archaea of the class Halobacteria [76, 77] and mainly display a head-tail morphology [78]. Previous work has suggested that the host range expansion of arTVs belonging to the *Hafunaviridae* family is primarily determined by mutations in the tail adhesin gene [75]. However, it is not known what role the antiviral defense systems play in determining the host range. To gain insight into the correlation between defense systems and viral virulence in haloarchaea, defense systems in 20 recently sequenced haloarchaeal genomes were analyzed [79] and compared with all complete Halobacteria genome sequences available in the RefSeq database (253 in total; downloaded in June 2025) using the PADLOC tool. These 20 strains were included in the analysis because their viral susceptibility to various arTVs is known [75], enabling their use in experimental work.

In total, 76 distinct system subtypes were identified in the genomes available in the RefSeq database (Figure 1, Supplementary Figure 1A and Table S1). The genome sequences contained between 0 and 23 defense systems, with an average of seven systems per genome (Figure 1A and Supplementary Figure 2A). The defense arsenal found among all genomes of the class Halobacteria in the RefSeq database showed that the most prevalent defense systems among members of the Halobacteria class are the putative defense candidates (PDC)-S70 (present in 83.5% of the analyzed genomes) and PDC-S27 (71.2%) (Figure 1B and Supplementary Figure 1A). PDC-S70 encodes a protein with a PIN nuclease domain, which is commonly associated with TA and Abi systems [74, 80]. PDC-S27, on the other hand, encodes a protein with an ATPase domain fused to members of the PD-(D/E)XK endonuclease superfamily, and is probably associated with nucleic acid targeting [74]. SoFic (51.3%), CRISPR-Cas type IB (33.1%) and Hma-embedded candidates (HEC)-06 (22.9%), HEC-05 (20.8%) were also found in high numbers in the haloarchaeal genomes (Figure 1B and Supplementary Figure 1A). Nearly every genome in the dataset encoded at least one RM system, including types I, II, III, and IV. The most prevalent subtypes were type II (33.5%) and type I (25%).

**Figure 1.**
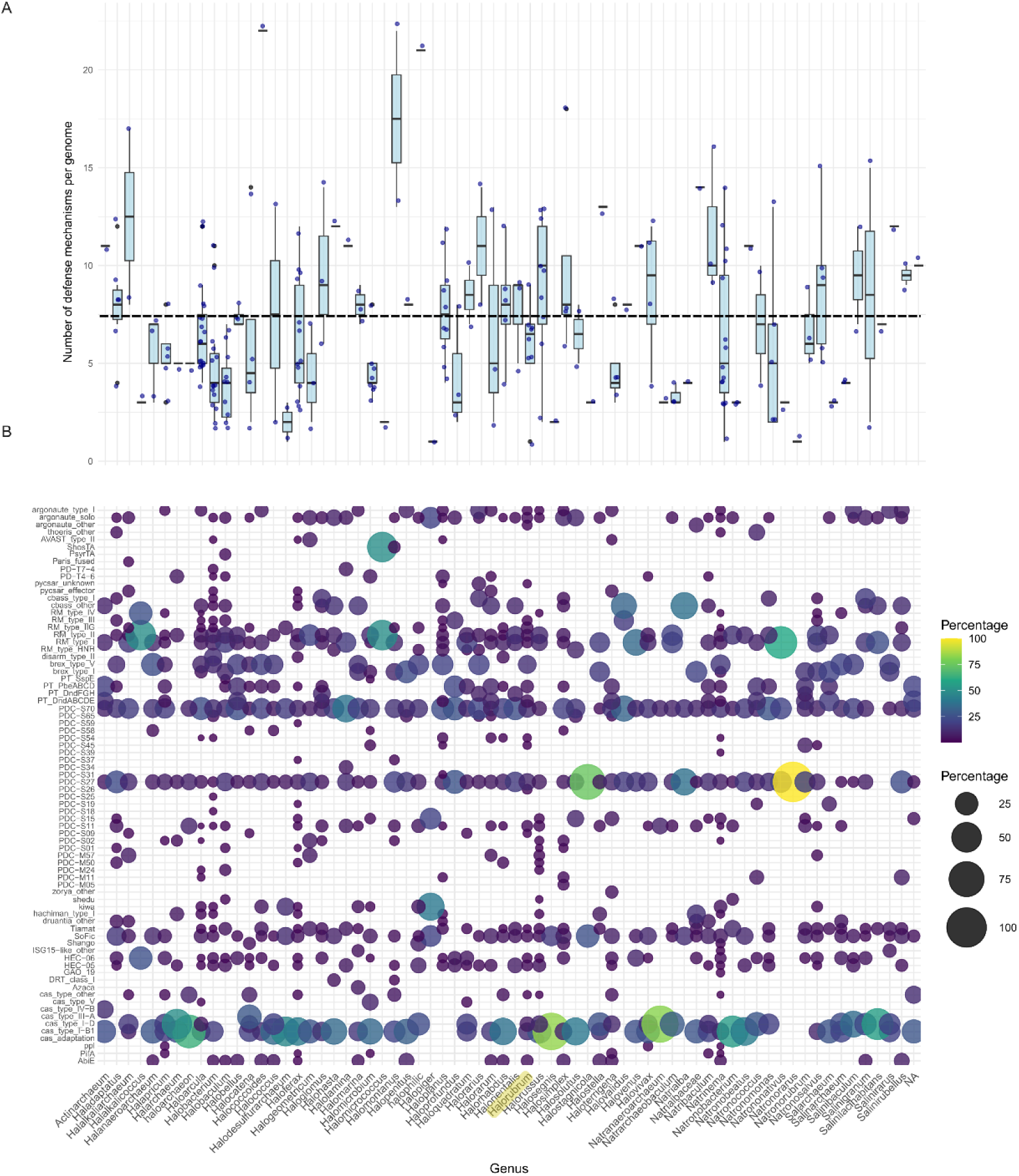
Defense systems found among all complete genome sequences of the class Halobacteria in the RefSeq database (n=253). (A) Number of defense systems per genus. Genera are listed in alphabetical order, and defense systems are ordered by mechanistic similarity. Each dot represents one strain. The average number of defense systems is represented as a black horizontal line. (B) Bubble plot showing the distribution of defense systems in each genus of Halobacteria. Dots represent the percentage of defense systems in each genus. The size and color of the dots are proportional to the percentage of the defense mechanism in that genus. The x-axis is the same for panels A and B. The genus *Halorubrum* is marked yellow on the x-axis.

Among the 20 recently sequenced haloarchaeal genomes used in this study, a total of 28 known defense system subtypes were identified, of which six were PDCs (Figure 2 and Supplementary Figure 1B). The number of defense systems per genome ranged from 0 to 13, with an average of six systems per genome (Supplementary Figure 2B). The most prevalent defense systems in these genomes are PDC-S70 and PDC-S27 (both present in 57.9% of the analyzed genomes). HEC-06 was present in 31.6% of the genomes followed by type II RM system and SoFic (both 26.3%). The gene locations of the identified systems are shown in Table S1 (Halobacteria) and Table S2 (new Haloarchaea genomes of this study).

**Figure 2.**
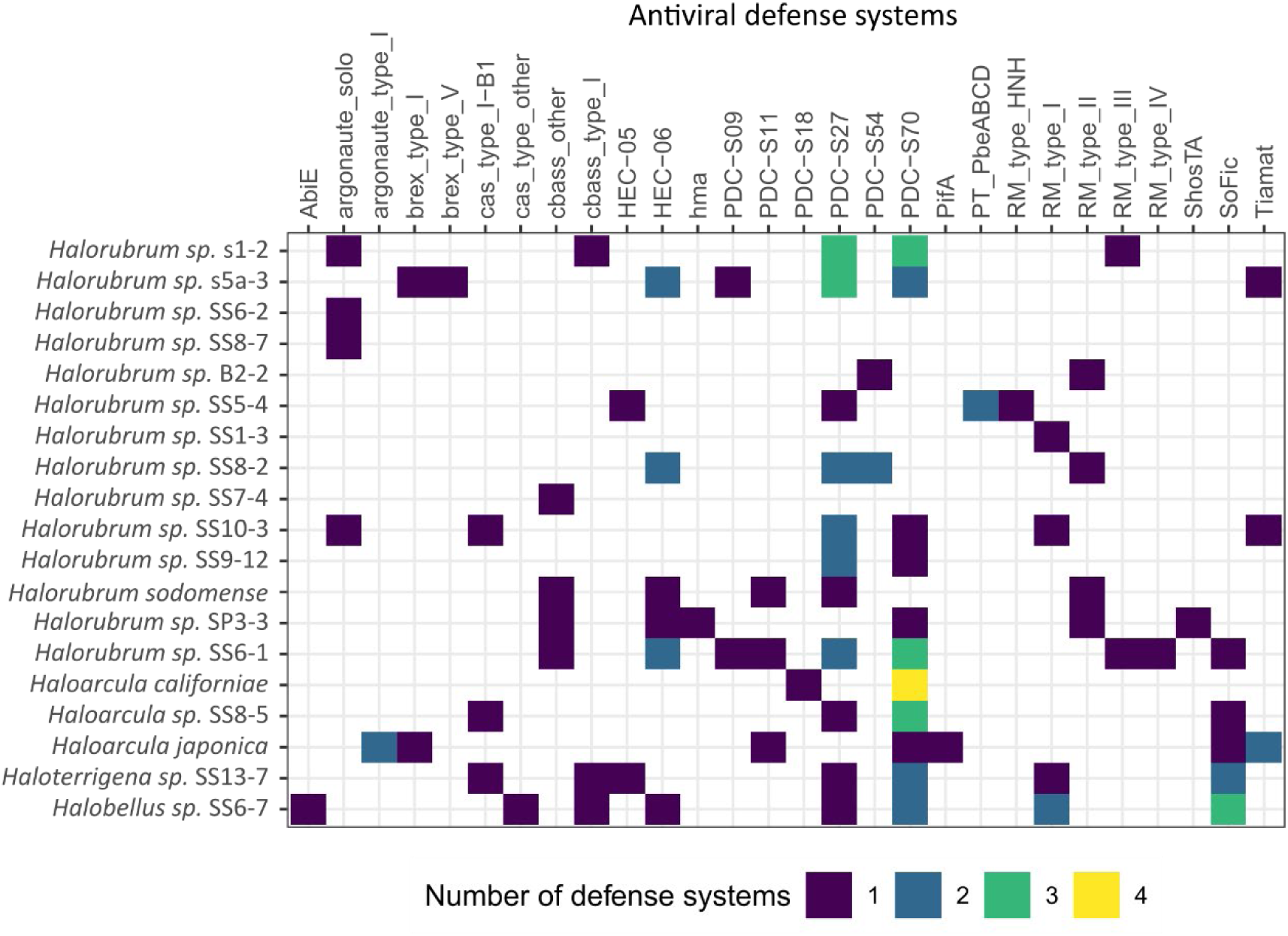
Types and number of defense systems identified in the set of 20 recently sequenced haloarchaeal genomes used in this study. The number of defense systems is indicated in purple, blue, green or yellow for 1, 2, 3 or 4, respectively. in *Halorubrum* sp. SS10-9 is not shown, as no defense systems were identified.

A comparison of the occurrence of defense systems between the 20 haloarchaeal sequences from this study and the RefSeq dataset revealed a similar distribution with some deviations (Supplementary Figure 1). Although all identified systems were also present in the RefSeq dataset, some defense systems were underrepresented among the 20 newly sequenced haloarcheal genomes. Notably, SoFic was present in only 26.3% of the genomes, compared to 51.3% in the Halobacteria RefSeq dataset. HEC-05 was present in only 10.5% of the genomes (20.8% in Halobacteria RefSeq) and a similar trend was seen for CRISPR-Cas type IB (20.1% vs 33.1% in the Halobacteria RefSeq). HEC-06 was more prevalent in our subgroup than in the RefSeq dataset (31.6% vs 22.9% in the Halobacteria RefSeq). Multiple copies of the same system were found in some genomes. For example, four copies of the PDC-S70 system were found in the genome of *Haloarcula californiae* representing the highest number observed in our dataset (Figure 2).

Overall, these results show that haloarchaea encode a diverse array of defense systems, which vary in their distribution and abundance across genomes. The distribution and abundance of potential defense systems in the experimental set of 20 strains closely mirrored those observed in haloarchaeal strains in general, indicating that the selected set of strains are representative of all haloarchaea.

### Relationship between antiviral defense systems and resistance to viral infection

To investigate the effect of defense systems on the susceptibility of haloarchaea to arTV infection, ten archaeal viruses belonging to the class *Caudoviricetes* were selected. Their infectivity on the 20 newly sequenced strains was determined in a previous study and expanded in the current study [75] (Figure 3A). Virus susceptibility of each haloarchaeal strain was determined based on plaque formation by cross-testing all viruses against all strains. Of 200 virus-host cross-tests (10 viruses on 20 hosts), 80 virus-host pairs showed positive infection, while 120 cross-tests did not lead to plaque formation (Figure 3A). To determine whether the failure of the viral infection cycle was due to the absence of a receptor (halting the infection cycle at the cell surface) or functioning of the intracellular defense systems (halting the infection at the intracellular stage), adsorption assays were performed to analyze the binding of non-plaque-forming viruses to the cell surface. Results showed that only a small number of the non-infectious viruses were able to adsorb on the cells (17 out of 120 cross-tests) (Figure 3A). To evaluate whether a higher number of defense systems correlates with increased virus resistance, a linear regression analysis was performed. The analysis revealed a weak negative correlation between the number of defense systems and resistance to viral infection (r2= 0.23 and P=0.06 Figure 3B). Only samples with adsorption levels statistically different from the cell-free control were considered positive.

**Figure 3.**
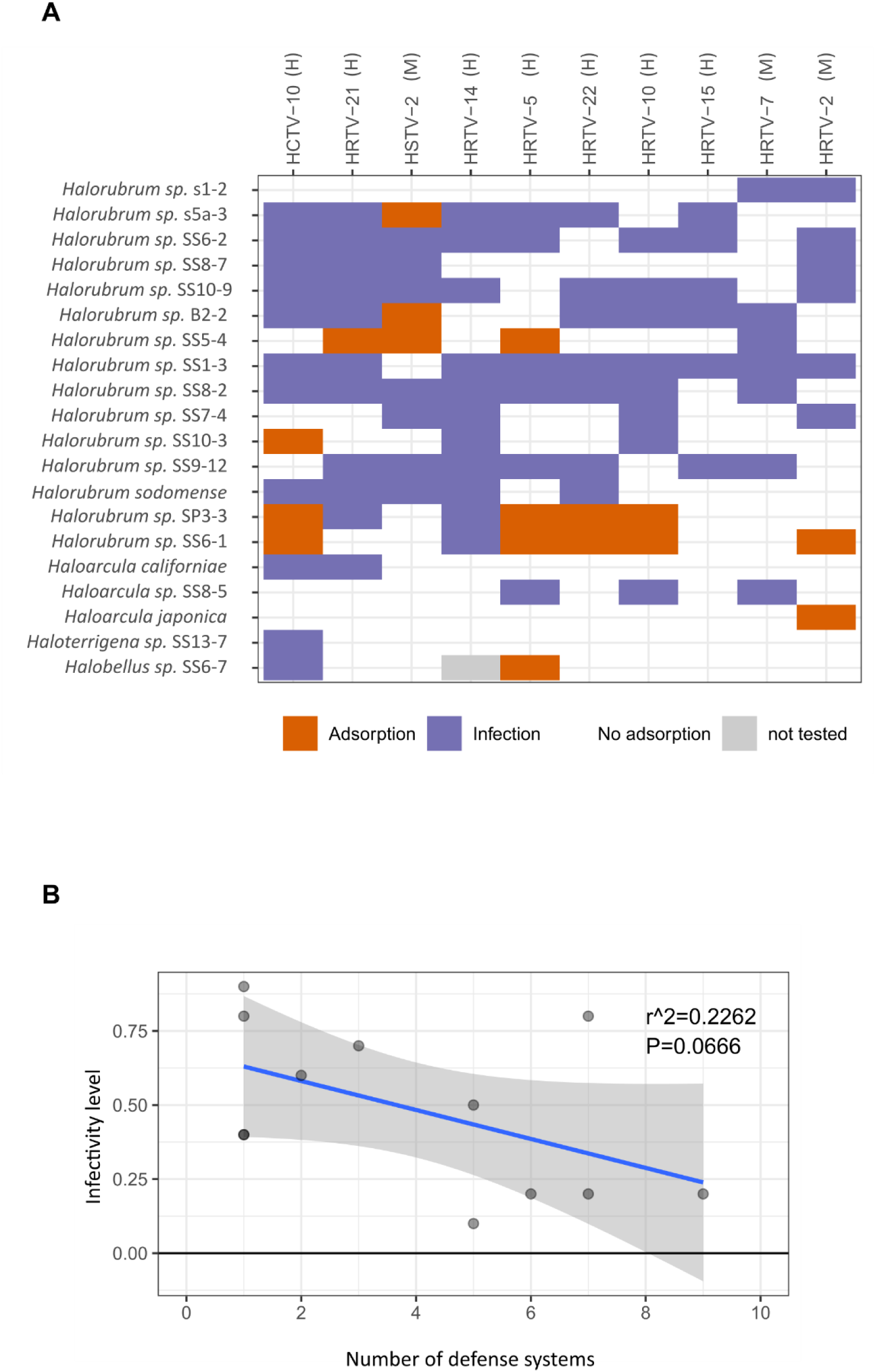
Presence of antiviral defense systems and viral susceptibility (A) Infectivity and binding of arTVs on 20 haloarchaeal strains. Haloarchaeal strains are ordered based on Average Nucleotide Identity (ANI) while viruses are grouped according to the classification of their tail fiber adhesin proteins, as defined by [75]. Virus-host interactions are depicted as infection (plaques formed; blue), adsorption but not infection (red), or no interaction (white). All viruses belong to the *Hafunaviridae* family and their genus is indicated next to their names; H, *Haloferacalesvirus*; M, *Mincapvirus*. (B) Linear regression analysis assessing the correlation between the number of defense systems in the host strain genome and the level of virus resistance.

Two strains with a relatively broad intracellular resistance against viruses were identified: *Halorubrum* sp. SS6-1 (against five viruses) and *Halorubrum* sp. SP3-3 (against four viruses). HCTV10, HRTV5, HRTV-22, HRTV-10 and HRTV-2 viruses adsorbed to SS6-1 and HCTV10, HRTV5, HRTV-22, and HRTV-10 adsorbed to SP3-1 but were unable to produce successful progeny. The genomes of *Halorubrum* sp. SS6-1 contained 13 defense systems, exceeding the average of six per genome observed across the dataset (Figure 2, Supplementary Figure 2A). In comparison, *Halorubrum* sp. SP3-3 encoded six defense systems. Defense systems detected in these strains included CBASS, HEC-06, ShosTA, SoFic, various PDCs and restriction-modification systems (Figure 2). Another strain showing intracellular resistance to multiple viruses was *Halorubrum* sp. SS5-4. It showed intracellular resistance to three viruses (HRTV-21, HSTV-2 and HRTV-5) and its genome harbored five defense systems.

Other strains were highly susceptible to viral infection, as they were infected by at least 8 of the 10 tested viruses. These included *Halorubrum* sp. SS6-2, SS10-9, SS1-3, and SS8-2. All of these strains possessed either one or no identifiable defense systems in their genomes. Interestingly, the only outlier among the highly virus-susceptible strains was *Halorubrum* sp. SS8-2, which encodes seven defense systems and is susceptible to eight of the tested viruses. Thus, overall, this study found that strains with a limited number of defense systems generally were more susceptible to viral infection compared to strains with above average numbers of defense systems.

Immunity against individual viruses might be related to distinct antiviral defense systems. To evaluate the contribution of individual defense systems to virus immunity, the pairwise correlations were assessed between the presence of specific defense systems and resistance to virus infection in virus-host pairs that showed adsorption but no successful infection (indicated by the absence of plaques) (Figure 4). While most correlations were weak, likely due to the relatively small sample size, a few strong correlations were observed (>0.75). Notably, the RM type HNH and the PT_PbeABCD system appear to confer protection against HRTV-21 infection. Among the systems identified, ShosTA, Type IV RM and hma displayed the broadest protection activity, primarily targeting HRTV-22 and HRTV-10 infections. In contrast, PDC-S11 and argonaute type I showed protection only against HRTV-2, while HEC-05 showed specificity against HRTV-21 infection. These results suggest that these defense systems may play targeted roles in antiviral protection of these archaeal strains.

**Figure 4.**
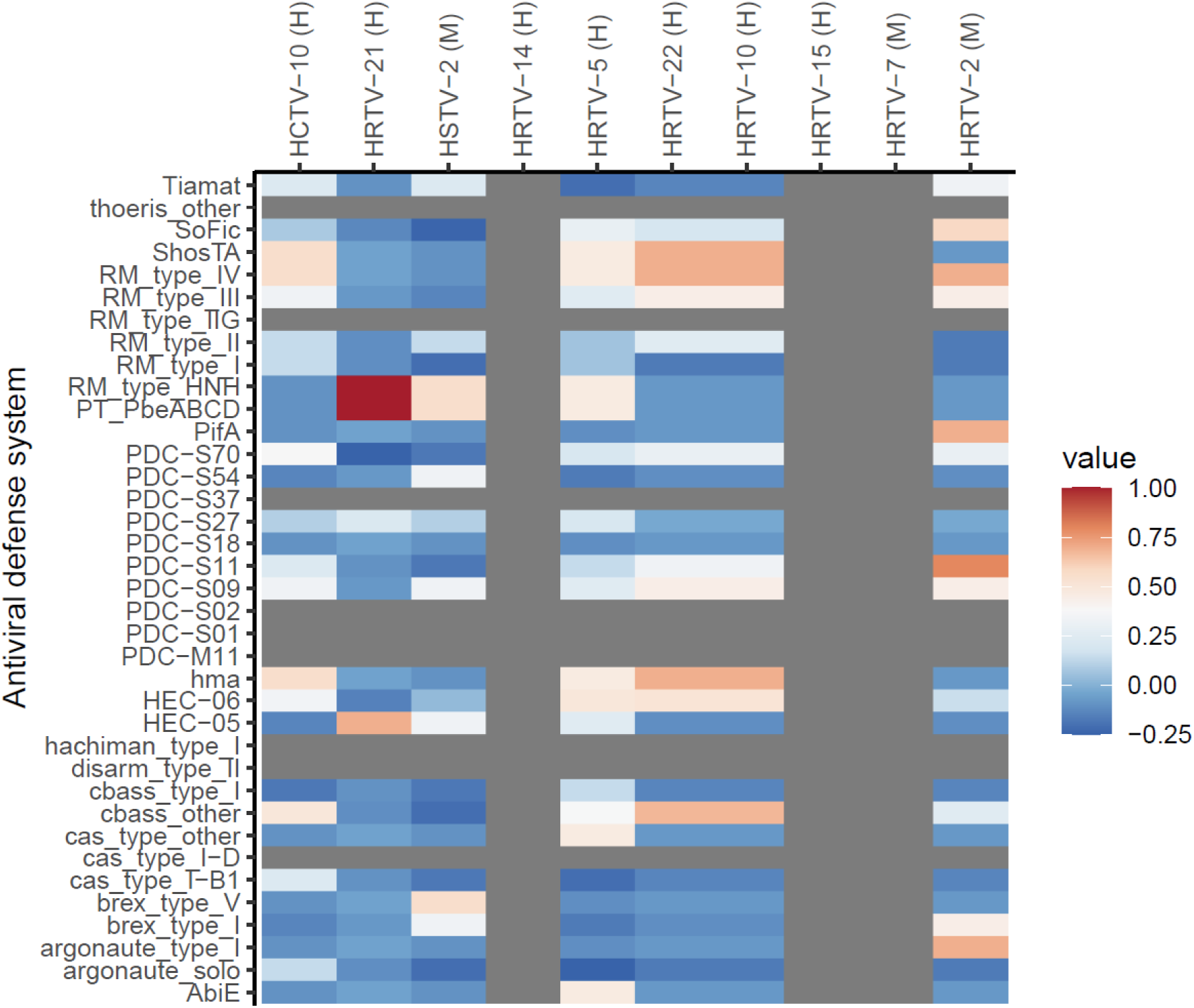
Correlation matrix between the antiviral defense systems encoded in haloarchaea and infectivity of arTVs. The color scale (bar on the right) represents the magnitude of the phi-coefficient, indicating the correlation level between the presence of a specific defense system and resistance to a certain virus. Higher phi-coefficient values indicate a stronger association between the presence of the defense system and the immunity against the virus. Grey values indicate that no virus-defense system occurrence appeared in our dataset. All viruses belonged to the *Hafunaviridae* family and their genus is indicated next to their names; H, *Haloferacalesvirus*; M, *Mincapvirus*

### Viruses might overcome archaeal defense systems via diverse anti-defense proteins

Besides the numerous antiviral defense systems identified in bacteria and archaea, over the last years, a variety of viral counter-defense strategies have also been discovered. These mechanisms help viruses to circumvent host defense. Examples of such counter-defense strategies include genomic modifications, either by mutations that alter the recognition sites targeted by host defense systems or via base modifications [81–83], the formation of nucleus-like compartments that shield viral genome from host nucleases [84], the protection by anti-CRISPR (Acr) proteins to inhibit CRISPR-Cas activity [85], and the use of accessory genes which likely interface with host processes during infection [86].

To explore the potential impact of viral counter-defense systems on host susceptibility, our analysis focused on systems with experimentally validated mechanisms of action. Among those identified, RM type II, SoFic, and CRISPRCas were the most prevalent in our dataset. Viruses have developed different systems to escape restriction by host RM systems, including the production of antirestriction proteins, avoidance of recognition sequences in their genomes, and base modifications [87–91]. To investigate this, the known recognition sequences of restriction modification enzymes encoded in haloarchaea genomes were retrieved from the REBASE database and their frequencies in the viral genomes were calculated (Table S3). All viruses lack the CTCGAG, GAGCAGC, GATC, GCATGC, GTCGAC, GWGCWC, and HGCWGCK sequences, whereas CCWGG was present only in HSTV-2 and CTAG in HRTV-22. Moreover, six of the 10 viruses analyzed encode a type IIG restriction-modification enzyme (Figure 5). This suggests that these viruses may overcome host RM systems either by encoding their own methyltransferases or by avoiding specific motifs in their genomes [75, 92, 93]. The SoFic system is composed of a stand-alone protein with a Fic domain, which is a protein AMPylase, and is considered as a member of toxin-antitoxin system group [94]. Other TA defense systems identified include ShosTA and PifA. Notably, many of the viruses analyzed in this study, encode a DUF4258 domain-containing protein (IPR025354) (Figure 5). Structural modeling indicates that proteins in this family adopt a fold characteristic of bacterial toxins, suggesting it functions as the toxin component in a TA system [95], which could function as a counter-defense to overcome toxin-antitoxin systems in the hosts. CRISPR-Cas did not appear to play a significant role in antiviral defense in our data set (Figure 4). However, anti-CRISPR (Acr) proteins that block CRISPR-Cas responses have been identified in archaeal viruses [96, 97]. Several of the viruses analyzed in this study encode putative Cas4 nucleases (Figure 5), which may function as Acr proteins [98–100]. Thus, counter-defense systems are encoded by the viruses used in this study. This may impact their host range and thus the correlation between host range and anti-viral defence systems analyzed in this study.

**Figure 5.**
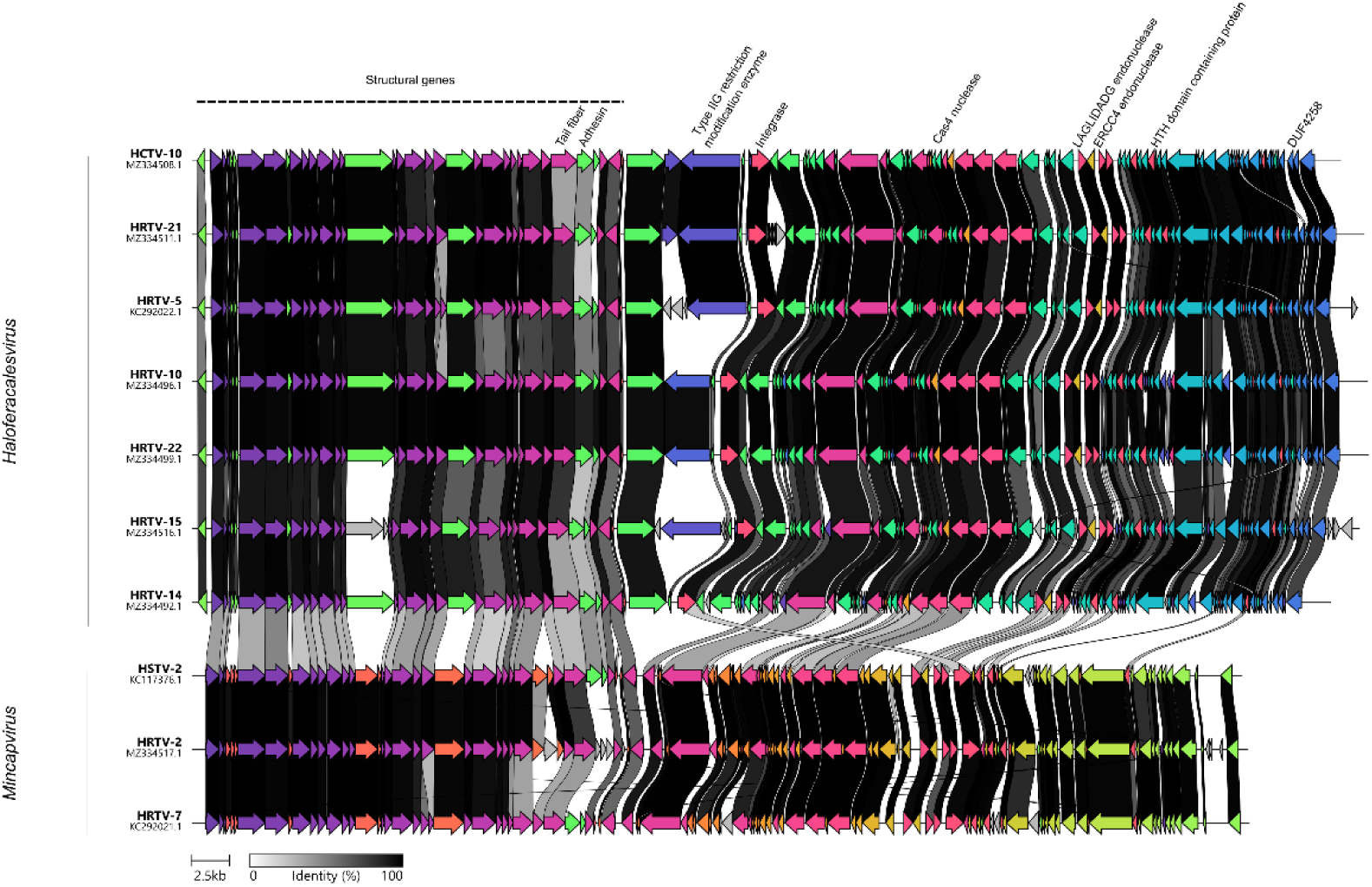
Comparative genome alignment of arTVs from the family *Hafunaviridae* (*Haloferacalesvirus* and *Mincapvirus* genera as indicated on the left side) used in this study. Virus names and accession numbers are shown on left. Coding sequences are represented by arrows. Arrow colors represent gene clusters identified by Clinker. Grey bars between the genomes represent homologous genomic regions and the intensity represents the level of nucleotide identity. Location of the structural genes are shown, with special emphasis on the tail fiber and adhesin, which have been proposed to be one of the main host range determinants. Other predicted gene functions, include counter-defence systems, such as type IIG restriction modification enzymes (counter defense against RM systems), Cas4 nuclease (against CRISPR-Cas), LAGLIDADG and ERCC4 endonucleases (DNA cleavage), HTH domain containing protein (for controlling gene expression) and DUF4268 domain protein (counter defense against TA). Putative integrase proteins are indicated. The alignment was created using Clinker with default settings.

## Discussion

The prevalence of defense systems was analyzed in a representative set of haloarchaeal strains to examine their relationship to virus host range. The set of 20 recently sequenced haloarchaeal strains [79] exhibited frequencies of genomically encoded defense systems comparable to those observed across the broader Halobacteria class (Supplementary Figure 1). The most prevalent defense systems were PDC-S70 and PDCS27, which is in line with a recent analysis of the occurrence of different defense systems among archaea [74]. Interestingly, individual haloarchaeal genomes often encoded multiple distinct defense systems (Figure 2). Some of the strains also showed redundant systems in their genomes (Figure 2). The co-occurrence of multiple defense systems in a single genome indicates that, like in bacteria, archaeal defense systems might act synergistically against viral infections [101].

Notably, among the tested viruses, only HCTV-10, HRTV-21, HRTV-5, HRTV-10, and HRTV-7 were capable of infecting strains outside the *Halorubrum* genus (Figure 3A). This is in line with previous studies that suggest that the host range of viruses belonging to the *Hafunaviridae* family is restricted only to specific species in a limited number of genera [34, 102, 103].

The virus-host pairs that showed no infection may have resulted from one of the following events: (i) failure of virus to adsorb to the host cell surface due to the lack of a suitable receptor, or (ii) inability to propagate due to an intracellular defense mechanism [35]. Adsorption assays of the non-plaque forming viruses revealed that only a small number of them were able to adsorb on the cell surface, which suggests that, for most viruses, infection is blocked at the adsorption stage (Figure 3A). Consistent with previous observations [75, 104] the binding of arTVs is host-specific, as most of *Halorubrum* viruses were unable to infect or adsorb to the members of other genera such as *Halobellus* or *Haloarcula* (Figure 3A). Consistent with previous reports, no clear correlation between the host recognition and virus taxonomy was seen [75]. However, it should be noted that, due to the standardized adsorption conditions, some adsorption events may have been overlooked. In summary, the host range of haloarchaeal viruses seems to be determined for a large extend by the presence of viral receptors on the host cell surface. In addition, a weak correlation was found between the number of defence systems and viral host range.

This study did not reveal a clear correlation between particular individual defense systems and resistance to specific viruses in archaea. It is important to note that these findings may be somewhat limited, likely due to the relatively modest number of virus-host pairs analyzed. Moreover, the tools used to detect defense systems in haloarchaea are predominantly based on bacterial datasets. Previous studies have shown that some defense systems are exclusively present in archaea [74] and certain archaeal systems have demonstrated homology to eukaryotic immune systems [73]. Moreover, archaea may employ non-DNA-encoded defense systems, such as extracellular vesicle (EVs) secretion, which have been shown to inhibit phage infection in bacteria [105, 106]. As members from all three domains of life secrete EVs [107, 108], it is possible that EVs produced by haloarchaea also contribute to antiviral defense. Other potential mechanisms may include receptor masking [109, 110], mutations in receptor genes [111], production of extracellular matrices, or biofilms [112, 113]. Moreover, quorum sensing has been shown to regulate defense systems in bacteria [114–118]. Future identification of archaeal specific anti-viral defense systems may strengthen the observed correlations between viral susceptibility and anti-viral defense systems in archaea. Together, these findings highlight the importance of exploring defense strategies in archaea, paving the way towards a deeper understanding and characterization of defense systems in haloarchaea.

While much remains to be discovered, our findings indicate that resistance typically occurs at the stage of adsorption, where the viral receptor-binding proteins interact with specific cellular receptors. When absorption is successful, haloarchaea rely on a variety of intracellular defense systems, such as RM, CRISPR-Cas and TA systems, to combat virus infection. A weak correlation was observed between the number of encoded defense systems by a strain and its susceptibility to viral infection, suggesting that strains with a greater number of antiviral defense systems are generally less susceptible. However, it is likely that additional uncharacterized defense systems that escaped our bioinformatic analysis also contribute to antiviral protection of archaeal cells and may make the correlation stronger in future studies.

Overall, the results of this study show a similar trend as reported earlier for this relationship in bacteria [35]. In the model archaeal group Haloarchaea, susceptibility to viruses is largely determined by the presence of viral receptors on the host cell surface, whereas in bacteria the defense systems play a much larger role. In addition, intracellular antiviral defense systems play a modest role in determining the viral susceptibility, as a higher number of defense systems generally correlates with increased viral resistance in the haloarchaeal strains analyzed. These findings offer a stepping stone for further analysis of host-range determinants of archaeal viruses and the role of antiviral defense mechanisms.

## Materials and Methods

### Strains and media

Haloarchaeal strains and viruses used in this study are listed in Tables S4 and S5. The modified growth medium(MGM) was prepared by diluting a 30% (w/v) stock of artificial salt water (SW) containing (per liter) 240 g of NaCl, 30 g of MgSO_4_·7 H_2_O, 35 g of MgCl_2_·6 H_2_O, 7 g of KCl, 80mM Tris HCl (pH 7.2) and 5mM CaCl_2_. Strains and viruses were grown aerobically at 37 °C in MGM liquid media containing 23% SW [119]. The MGM soft agar media was prepared with 4 g/L of agar and 18% SW, where MGM plates contained 14 g/L agar and 20% MGM.

### Virus stock preparation

Virus stocks were prepared using the double-layer plaque assay method. Cells were grown in liquid media and 300 μl of the culture (dense growth) and 100 μl of the appropriate virus dilution were mixed with 3 mL of melted soft agar and poured into a plate. After incubation (2-3 days), the soft agar from plates that showed semi-confluent plaque amounts was collected in a flask, and 2 mL of MGM liquid per plate was added. The mixture was incubated at 37 °C for 1.5 h with aeration. Cell debris was removed from the culture by centrifugation (11.000×g, 25 min, 4 °C).

### Adsorption assays

Cells in the mid-logarithmic phase (10^8^ CFU/mL) were mixed with viruses at a multiplicity of infection (MOI) of 10^-3^ and incubated for 1 h at 37 °C with aeration. Samples were taken and diluted 100-fold with ice-cold liquid medium. The cells were removed by centrifugation at 13.000 rpm for 2min at 4°C (table-top centrifuge), and the supernatant was used to determine the number of unbound viruses by double-layer plaque assay. In the control, viruses were mixed with liquid medium (no cells) to determine the total virus concentration. The number of adsorbed viruses was determined by subtracting the amount of non-adsorbed viruses from the total virus concentration. Samples exhibiting significant absorption compared to control values were determined with edgeR v4.0.16 [120], using exact tests with global dispersion estimates at p <0.05. Only samples in which the amount of unbound virus particles was statistically significant from the control were considered as positive adsorption. All adsorption assays were carried out in triplicate.

### Virus infectivity

The virus infectivity on different host strains was determined as described by Liu et al [75]. Briefly, 10 μl drops of undiluted and 100-fold diluted virus stocks were spotted on a soft-agar lawn of the hosts (double-layer method) and incubated at 37 °C. Plates were checked after 2 to 3 days for cell lysis and those host virus pairs showing host cell growth inhibition were further analyzed by double-layer plaque assay.

### Analysis of antiviral defense systems in haloarchaea genomes

The accessions and metadata of all complete Halobacteria genomes from RefSeq (accessed in June 2025) were retrieved, resulting in 253 sequences. The genome sequences of the 20 haloarchaea used also in the experiments were obtained in [79]. PADLOC v2.0.0 [121] was used with default settings to identify known antiviral defense systems in the genomes. Defense systems belonging to the DNA modification systems category were discarded as they are potentially incomplete systems. Analyses and visualization were done using a self-written R script (R version 4.4.1 (2024-06-14)), built from common packages (dplyr, tidyr and ggplot2). The correlation of defense systems and frequency of infection was assessed using the Pearson coefficient on logit-transformed frequency values.

### Analysis of viral genomes

Viral genomes were downloaded from NCBI (accessed in June 2025) (Table S5) and alignments of genome similarity were generated using clinker [122]. Annotations were done based on the already annotated genomes in NCBI.

## Acknowledgments

T.E.F.Q and Z.A.S were supported by funding from the Hector Fellow Academy. Further financial support is acknowledged from an ERC starting grant (101039446 ARCVIR) and a Vidi grant (Vidi.223.020) from the Netherlands Research Council (NWO) to T.E.F.Q. H.M.O. was supported by the University of Helsinki and the Research Council of Finland by funding for FINStruct and Instruct Centre FI, part of Biocenter Finland and Instruct-ERIC and Horizon MSCA 101120407. We thank Sari Korhonen for her skillful technical assistance at the University of Helsinki.

## Declaration of generative AI and AI-assisted technologies in the writing process

During the preparation of this work the authors used Microsoft Copilot in order to correct for grammatical accuracy and clarity. After using this tool, the authors reviewed and edited the content as needed and take full responsibility for the content of the publication.

## Supplementary

**Supplementary Figure 1.**
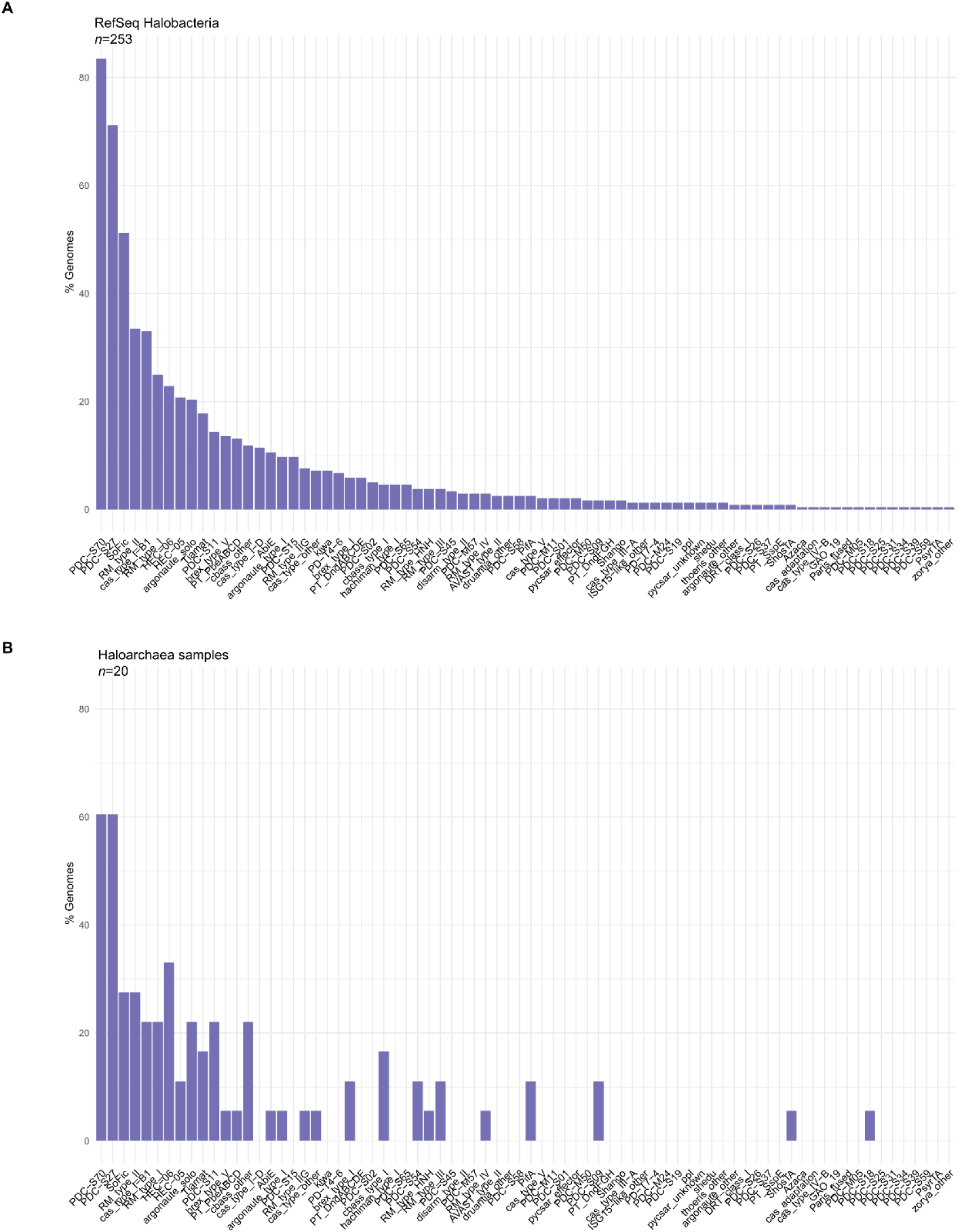
Diversity of defense systems found in (A) the genomes of 253 Halobacteria from the RefSeq database and (B) the genomes of the 20 strains in our collection [79], organized in a gradient from most (left) to least (right) abundant based on the RefSeq data in both panels A and B.

**Supplementary Figure 2.**
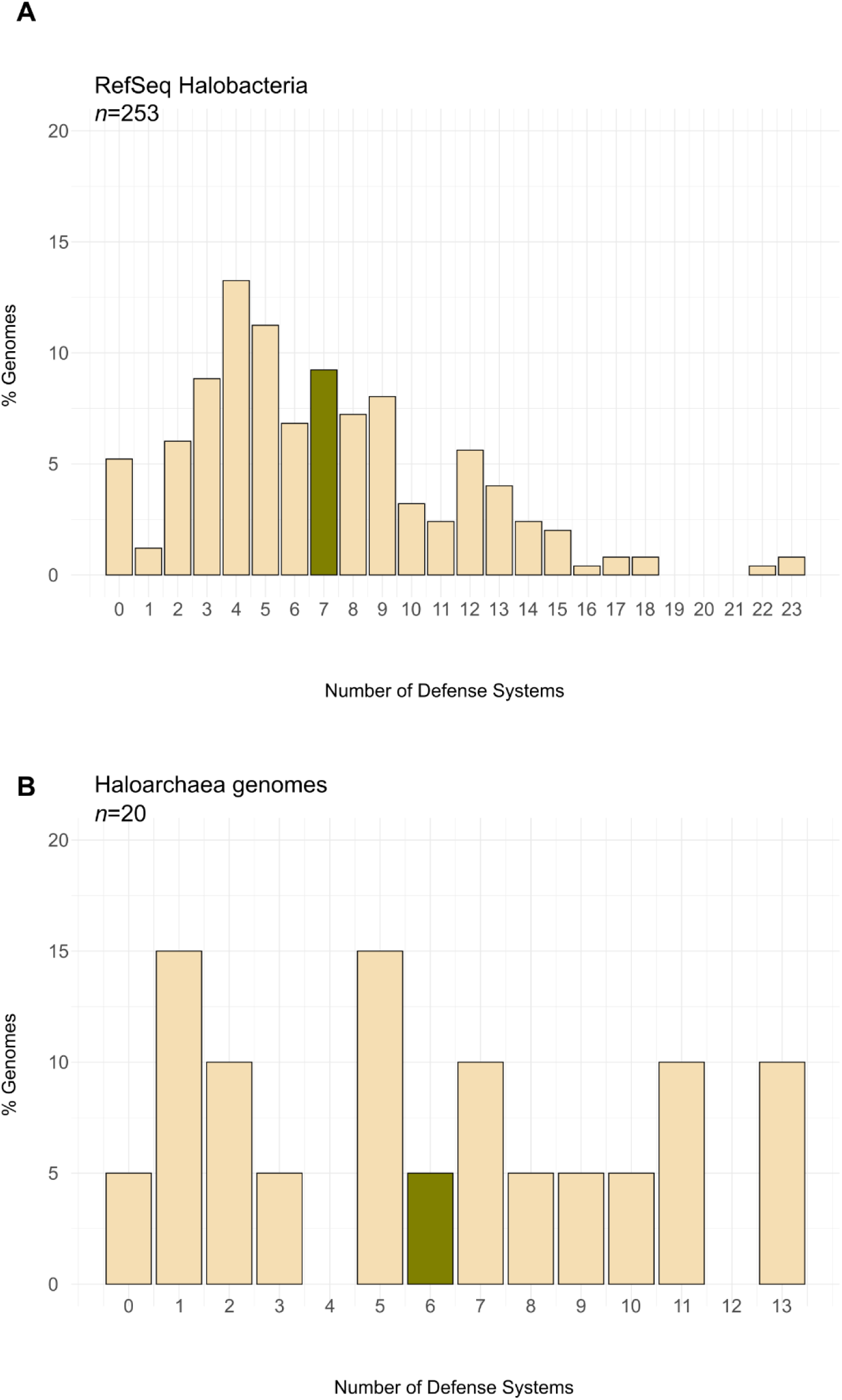
Number of defense systems per genome. (A) Number of defense systems per genome in Halobacteria from the RefSeq database (accessed in June 2025). (B) Number of defense systems per genome in 20 haloarchaeal strains from our collection. The average number of defense systems is shown in green.

**Supplementary Figure 3.**
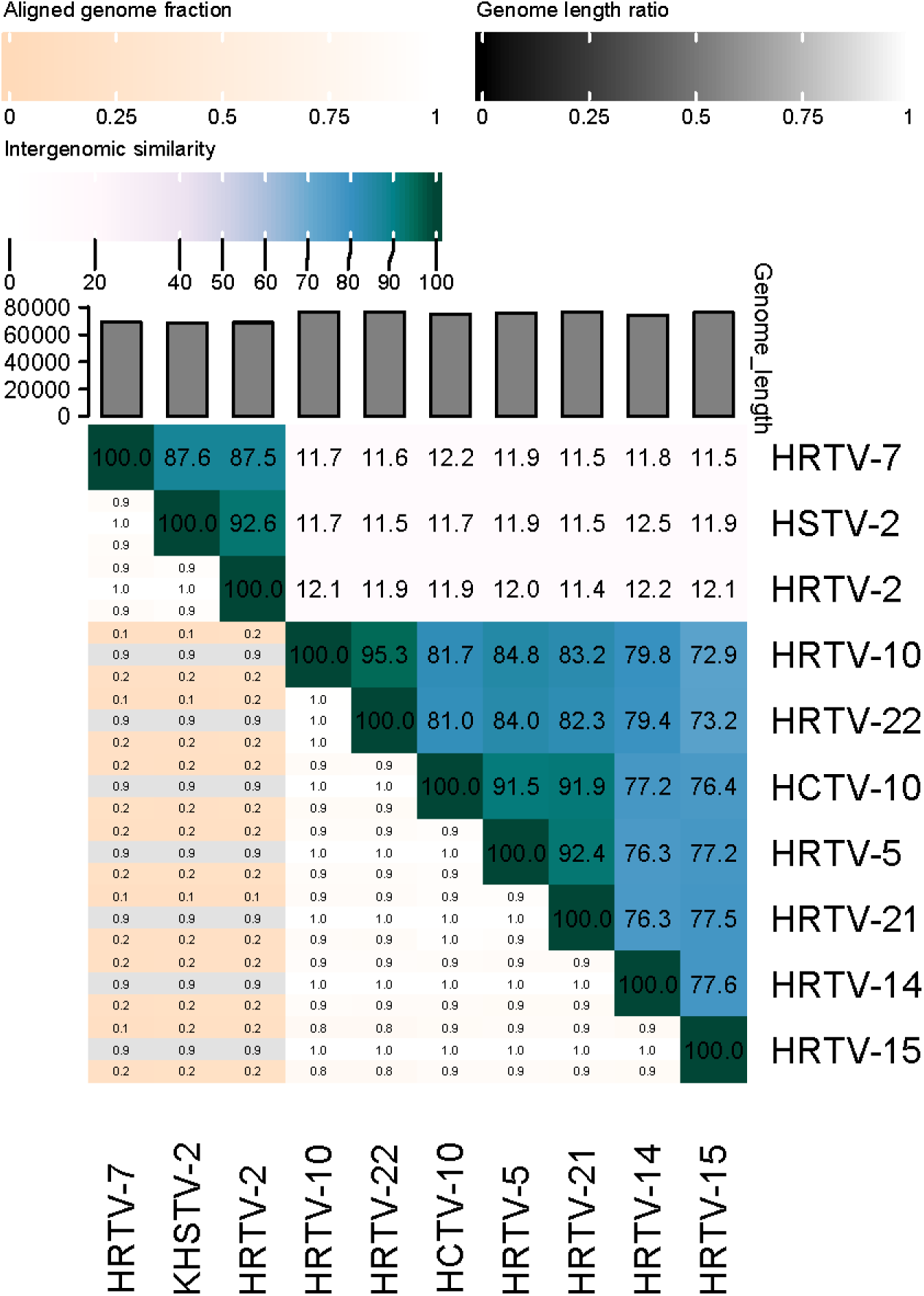
VIRIDIC heatmap showing the intergenomic similarity between viral genomes using a color gradient on the right hand. Darker colors represent higher intergenomic similarities. On the left side the three different values are shown; the fraction of the genomes that is aligned (top), the genome length ratio in each particular pair (middle) and the aligned genome fraction (bottom).

**Table S1**. PADLOC results showing the defense systems identified in the genomes of the complete Halobacteria genomes available in RefSeq (in June 2025).

**Table S2**. PADLOC results showing the defense systems identified in the genomes of the selected haloarchaea analysed in this study.

**Table S3**. Frequency of motifs detected by haloarchaeal RM systems in the genomes of viruses analyzed in this study.

**Table S4**. Archaeal strains used in this study.

**Table S5**. Archaeal viruses used in this study.

